# TEAD1 signaling modulates adrenal chromaffin cell maturation

**DOI:** 10.64898/2026.08.20.745987

**Authors:** Longqing Xia, Xue Liu, Fei Yan, Jingru Qu, Ying Zou, Mengrui Chai, Liying Zhu, Ruya Liu, Vijay K. Yechoor, Li Chen, Kai Zhang, Fuqiang Liu, Xinguo Hou, Feng Li

## Abstract

Chromaffin cells synthesize and secrete catecholamines to coordinate systemic stress responses and regulate diverse neuroendocrine and metabolic functions. However, the molecular mechanisms governing chromaffin-cell differentiation and their disruption in pheochromocytoma (PC) remain incompletely understood.

Here, through integrated analyses of human developmental atlases, patient-derived transcriptomic datasets, genetically engineered mouse models, and chromaffin organoids, we identify TEAD1 signaling as a critical regulator of chromaffin-cell differentiation and function. In vivo studies using a chromaffin cell-specific TEAD1 overexpression mouse model demonstrated that suppression of TEAD signaling markedly compromises chromaffin-cell differentiation and endocrine function. Additionally, compared with other TEAD family members, TEAD1 transcriptional activities are readily affected by sequences near the binding motif.

To identify therapeutically actionable regulators of TEAD1 signaling, we established a TEAD activity-based screening platform and identified the serotonin receptor HTR5A antagonist SB699551 as a potent modulator of chromaffin-cell state. SB699551 suppressed PC-cell proliferation in vivo, and remodeled catecholamines synthesis in primary human PC cells. Additionally, application of SB699551 to human PC tumor revealed a subpopulation of primary chromaffin cells sensitive to this compound. Mechanistically, CXXC5 and L1CAM were identified as downstream SB699551-TEAD1 signaling effectors mediating chromaffin-cell proliferation and differentiation.

Overall, we demonstrate that TEAD1 signaling is a fundamental mechanism regulating chromaffin cell differentiation and that modulation of TEAD1 signaling via SB699551 offers a new area of investigation in chromaffin cell biology.

## INTRODUCTION

Chromaffin cells are neuroendocrine cells located primarily in the adrenal medulla. These cells synthesize and release catecholamines, epinephrine and norepinephrine (NE), into the systemic circulation-a process essential for life through these transmitters’ coordination of the stress response. Moreover, catecholamine secretion by chromaffin cells is critical for a myriad of additional homeostatic processes especially throughout development and early life, including breathing initiation, regulation of heart rate and blood pressure, as well as the physiological response to hypoxia [1]. Catecholamines have marked metabolic effects, particularly on glucose metabolism [2], including increasing the production of glucose through glycogenolysis and gluconeogenesis in the liver [3]. Nevertheless, despite their functional importance, many fundamental questions remain concerning how chromaffin cells develop and maintain their differentiation.

Pheochromocytomas (PCs), catecholamine-secreting neuroendocrine tumors, arise from chromaffin cells. Due to the episodic release of excess catecholamines into the circulation, the classic symptoms of PC tumors include potentially life-threatening elevations in blood pressure, headaches, palpitations, anxiety, and diaphoresis [4]. PCs can also cause impaired glucose tolerance and hyperglycemia, which may lead to diabetes [5]. Since chromaffin cells remain functional in PCs, elucidating the mechanisms underlying their maturation and proliferation may pinpoint potential therapeutic targets for PCs.

The TEAD family of transcription factors serves as a central effector of multiple signaling pathways controlling cell fate determination, differentiation, and tissue homeostasis [6–7]. Among TEAD family members, TEAD1 has been implicated in the development and maintenance of β cells [8]. However, whether TEAD1 signaling regulates chromaffin-cell maturation and contributes to PC pathogenesis remains largely unknown. Moreover, most studies infer TEAD1 pathway activity based on TEAD1 expression levels, despite growing evidence that transcription factor abundance does not necessarily reflect transcriptional output [9–10]. In our previous studies, we found that TEAD1 can repress transcription by interfering with the binding of RNA polymerase II to chromatin, thereby resulting in reduced TEAD signaling activity [11]. However, whether this phenomenon occurs in vivo remained unclear.

Recent advances in single-cell transcriptomics have enabled comprehensive characterization of developmental trajectories and tumor-cell states at unprecedented resolution [12]. Nevertheless, the lack of functional approaches capable of directly monitoring TEAD1 signaling activity has limited our understanding of how TEAD1-dependent transcriptional programs shape chromaffin-cell identity. Furthermore, whether TEAD1 signaling can be exploited as a platform for therapeutic target discovery in PC has not been explored.

In this study, we integrated human developmental and tumor single-cell datasets to define the role of TEAD1 signaling during chromaffin-cell differentiation. Using a newly generated TEAD1 signaling reporter mouse, chromaffin-cell-specific TEAD1 gain-of-function models, and human pheochromocytoma samples, we demonstrate that TEAD1 transcriptional activity, rather than TEAD1 abundance, serves as a critical determinant of chromaffin-cell maturation. Mechanistically, we identify a two-stage model in which TEAD1 promotes early lineage progression, whereas optimal TEAD1 signaling activity is required for terminal chromaffin-cell maturation. Leveraging a TEAD1 activity-based screening platform, we further identify the serotonin receptor HTR5A as a therapeutically targetable regulator of TEAD1 signaling and define CXXC5 and L1CAM as downstream effectors mediating chromaffin-cell proliferation and maturation. Together, these findings establish TEAD1 activity as a functional biomarker of chromaffin-cell state and provide a framework for therapeutic target discovery in pheochromocytoma.

## RESULTS

### TEAD1 is selectively associated with chromaffin-lineage maturation during human adrenal development

To define the transcriptional regulators governing chromaffin-cell differentiation, we integrated and jointly analyzed 39 publicly available human embryonic and fetal single-cell RNA sequencing datasets spanning adrenal development. Developmental trajectory reconstruction resolved the major cellular populations along the chromaffin lineage, including neuromesodermal progenitors (NMPs), trunk neural crest cells (trunk NCs), Schwann cell precursors (SCPs), sympathoadrenal progenitors/bridge cells (SAPs), and mature chromaffin cells (CCs) (Fig.1A).

**Figure 1.**
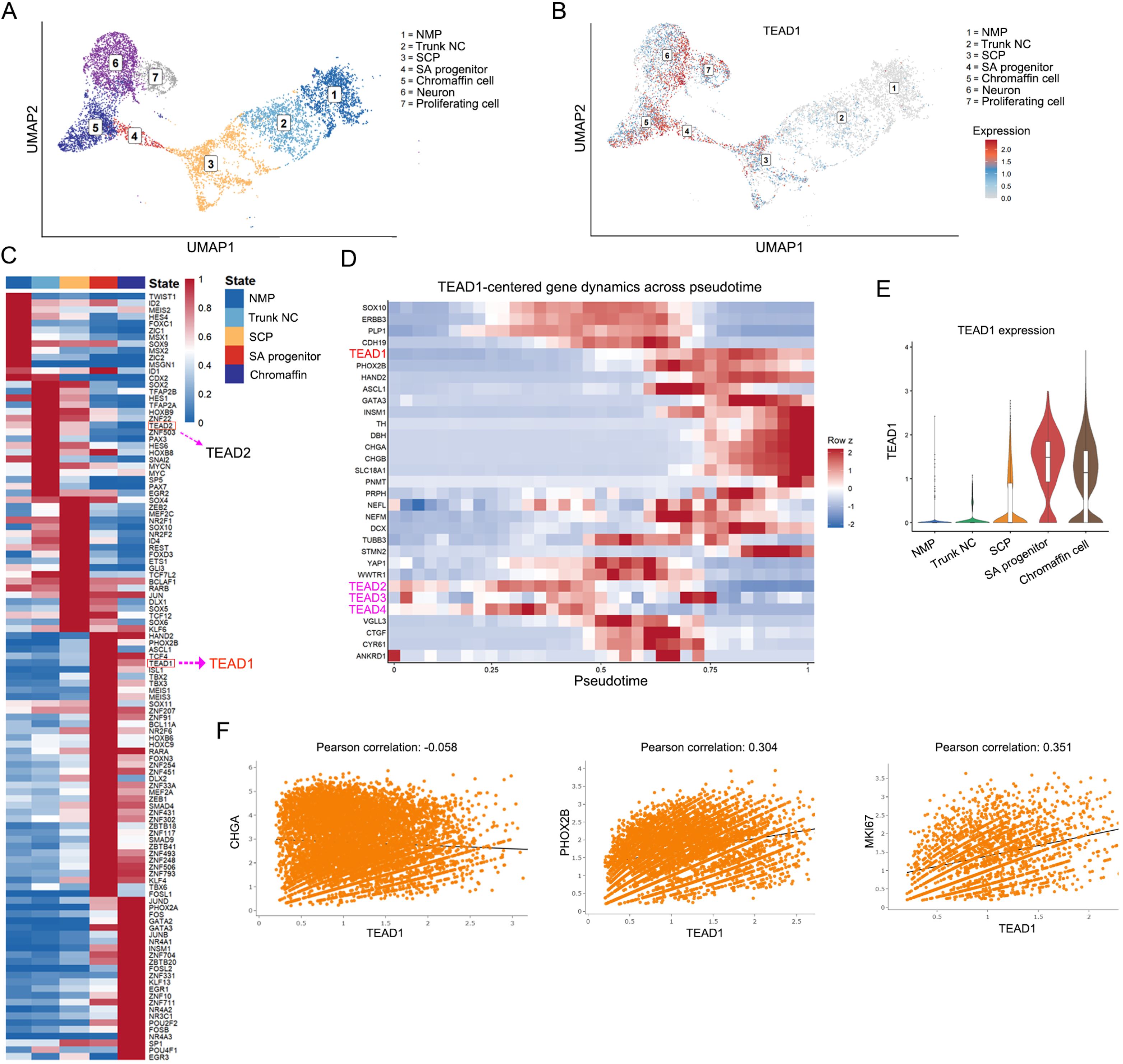
TEAD1 is associated with chromaffin lineage progression during human adrenal development. (A) UMAP visualization of integrated single-cell transcriptomic data from human embryonic and fetal adrenal tissues, identifying neuromesodermal progenitors (NMPs), trunk neural crest cells (trunk NCs), Schwann cell precursors (SCPs), sympathoadrenal progenitors/bridge cells (SAPs), and mature chromaffin cells (CCs). (B) Feature plot showing the distribution of *TEAD1* expression across developmental cell populations. (C) Regulon activity analysis of transcription factors across adrenal developmental lineages. (D) Pseudotime analysis of TEAD1 signaling-related genes during chromaffin-cell differentiation. (E) Violin plots showing *TEAD1* expression levels across distinct developmental cell populations. (F) Correlation analysis between *TEAD1* expression and *CHGA*, *PHOX2B*, and *MKI67* expression in embryonic chromaffin cells.

Feature plot analysis revealed a progressive increase in TEAD1 expression along the chromaffin developmental trajectory, indicating a strong lineage association (Fig.1B). To identify TEAD1-family members involved in chromaffin-cell specification, we performed regulon analysis and found that both TEAD1 and TEAD12 exhibited dynamic activity during lineage progression. Notably, TEAD1 regulon activity reached its highest level at the SAP stage and remained active during chromaffin-cell maturation, whereas TEAD12 activity was largely restricted to earlier developmental populations, including trunk NCs and SCPs, before declining sharply upon entry into the SAP and CC stages (Fig.1C). These findings suggest distinct developmental functions of TEAD1-family members, with TEAD1 emerging as the predominant TEAD1 factor associated with late-stage chromaffin differentiation.

Consistent with these observations, pseudotime analysis using Monocle demonstrated sustained TEAD1 engagement throughout chromaffin-cell maturation, whereas other TEAD1-family members displayed transient activity confined to early developmental states (Fig.1D). Thus, among all TEAD1-family factors, TEAD1 showed the strongest association with the acquisition and maintenance of chromaffin-cell identity.

To further characterize the biological context of TEAD1 expression, we performed correlation analysis within embryonic chromaffin cells. TEAD1 expression positively correlated with the chromaffin-lineage determinant PHOX2B and the proliferation marker MKI67, but showed no significant association with the pan-neuroendocrine marker CHGA (Fig.1E). These results suggest that TEAD1 expression is preferentially associated with proliferative and immature chromaffin-cell states during development, implicating TEAD1 in early lineage progression and expansion of chromaffin progenitors.

### TEAD1 expression is associated with a differentiated chromaffin-cell phenotype in pheochromocytoma

To investigate the role of TEAD1 in pheochromocytoma, we performed single-cell RNA sequencing on three pheochromocytoma samples and one adjacent adrenal tissue. Unsupervised clustering identified ten major cellular populations, including chromaffin cells (Fig. 2A, SupFig. 1A). Within the chromaffin-cell compartment, correlation analysis revealed a positive association between TEAD1 and the lineage-defining transcription factor PHOX2B (Fig. 2B), suggesting a potential link between TEAD1 expression and maintenance of chromaffin-cell identity.

**Figure 2.**
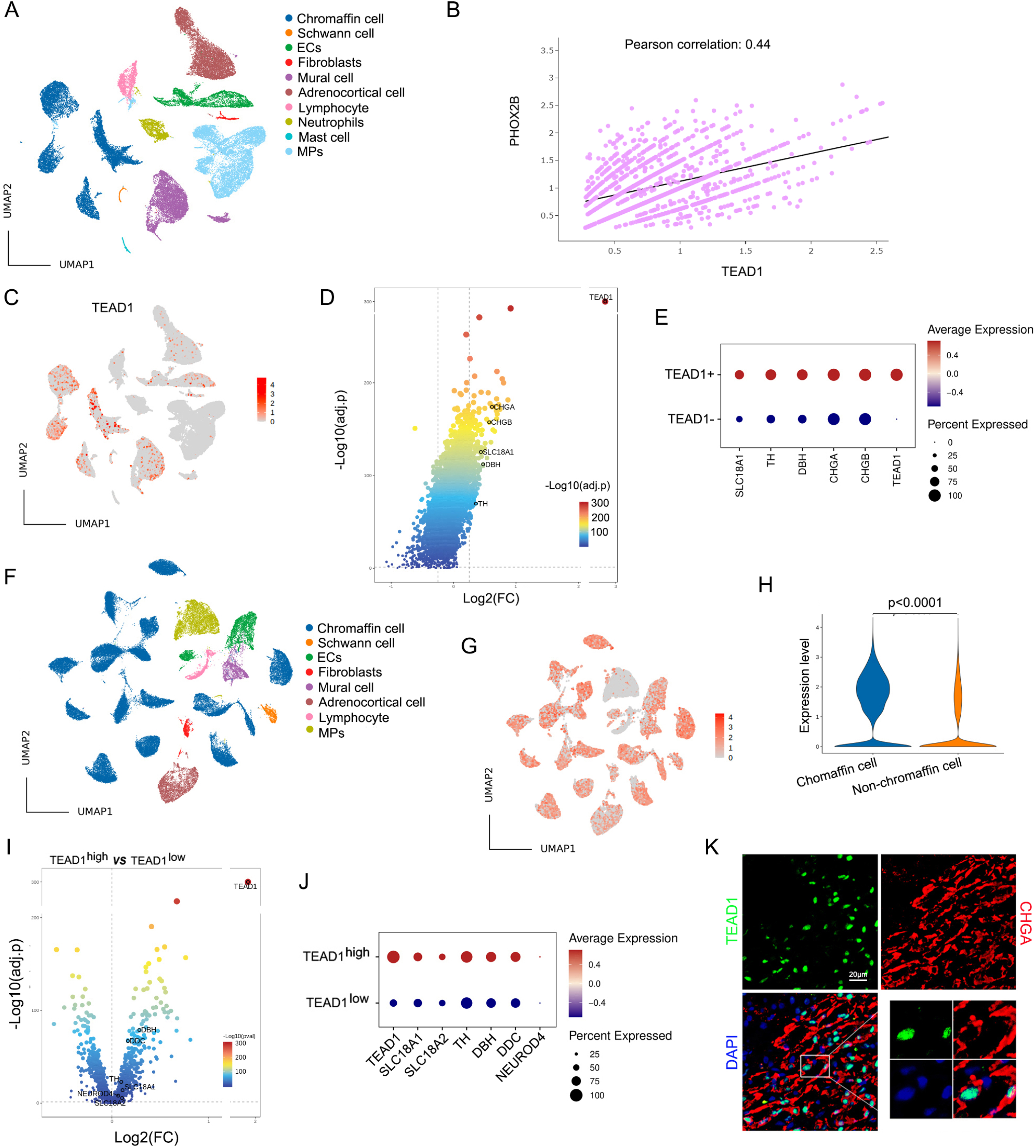
TEAD1 expression is associated with a differentiated chromaffin-cell state in human pheochromocytoma. (A) UMAP visualization of single-cell RNA sequencing data from three pheochromocytoma (PC) samples and one matched adjacent adrenal tissue, showing the major cellular populations identified in the tumor microenvironment.(B) Correlation analysis between *TEAD1* and *PHOX2B* expression in tumor chromaffin cells. (C) Feature plot showing the distribution of *TEAD1* expression across individual cell populations from PC and adjacent adrenal tissues. (D) Volcano plot showing differentially expressed genes between *TEAD1*-positive and *TEAD1*-negative chromaffin cells. (E) Bubble plot showing representative genes differentially expressed between *TEAD1*-positive and *TEAD1*-negative chromaffin cells. (F) UMAP visualization of single-nucleus RNA sequencing data from 18 human pheochromocytoma and normal adrenal samples, identifying chromaffin cells and other major adrenal cell populations. (G) Feature plot showing *TEAD1* expression across cell populations in the integrated single-nucleus dataset. (H) Violin plots comparing *TEAD1* expression levels between chromaffin cells and non-chromaffin cell populations. (I) Volcano plot showing differentially expressed genes between *TEAD1*^high^ and *TEAD1*^low^ chromaffin cells in the integrated pheochromocytoma cohort. (J) Bubble plot showing representative genes enriched in *TEAD1*^high^ and *TEAD1*^low^ chromaffin cells. (K) Representative immunofluorescence staining demonstrating TEAD1 expression in human pheochromocytoma specimens. Green, TEAD1. Red, CHGA. Blue, DAPI.

Feature plot analysis demonstrated that TEAD1 expression was highly enriched in chromaffin cells, whereas TEAD12, TEAD13, and TEAD14 were minimally expressed (Fig. 2C, SupFig. 1B). Differential expression analysis comparing TEAD1-positive and TEAD1-negative chromaffin cells revealed significant upregulation of key neuroendocrine and catecholaminergic markers, including CHGA, CHGB, SLC18A1, DBH, and TH, in the TEAD1-positive population (Fig. 2D-E). These findings suggest that TEAD1 expression is associated with a more differentiated chromaffin-cell state.

To validate these observations in an independent cohort, we reanalyzed a published single-nucleus RNA-sequencing dataset comprising 18 pheochromocytoma specimens (Fig. 2F). Consistent with our findings, TEAD1 was the predominant TEAD1-family member expressed in tumor chromaffin cells (Fig. 2G-H, SupFig. 1C). Stratification of chromaffin cells into TEAD1^high^ and TEAD1^low^ populations (SupFig. 1D) further demonstrated enrichment of catecholamine biosynthetic and secretory genes, including SLC18A1, SLC18A2, TH, DBH, and DDC, in TEAD1^high^ cells (Fig. 2I-J), supporting a strong association between TEAD1 expression and maintenance of the differentiated chromaffin-cell program.

Finally, immunofluorescence staining of an independent cohort of 14 human pheochromocytoma specimens detected TEAD1 protein expression in the majority of tumors (10/14), confirming the widespread presence of TEAD1 in human pheochromocytoma (Fig. 2K, SupFig. 2A-C).

### Excessive TEAD1 expression impairs terminal chromaffin-cell maturation through suppression of TEAD1 signaling activity

To determine the physiological role of TEAD1 in vivo, we generated chromaffin-cell-specific TEAD1 overexpression mice (cTOV, Fig. 3A). Unexpectedly, cold-stimulation experiments revealed a significant reduction in circulating epinephrine levels in cTOV mice (Fig. 3B), suggesting impaired chromaffin-cell function despite increased TEAD1 expression.

**Figure 3.**
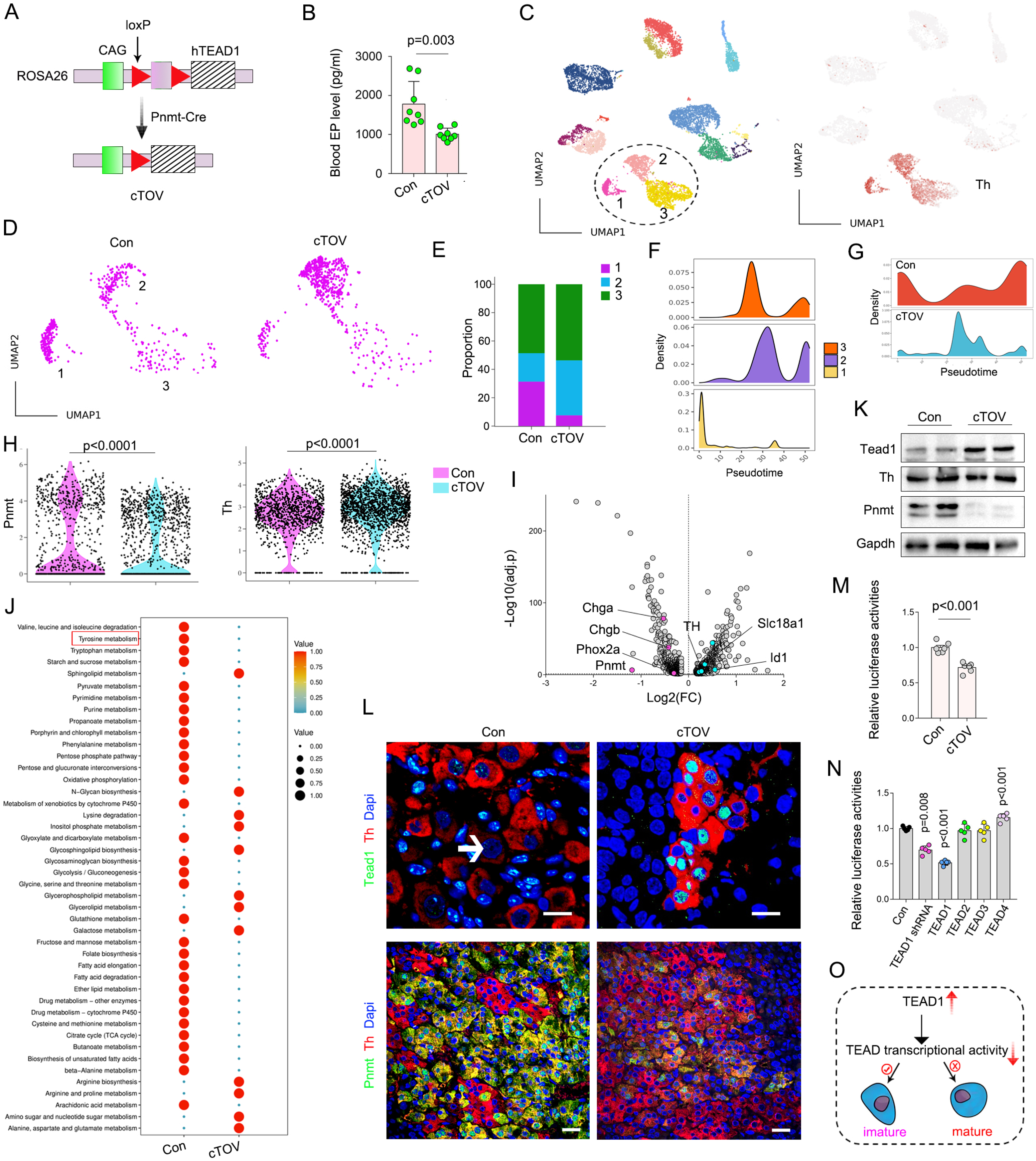
Excessive TEAD1 expression impairs terminal chromaffin-cell maturation through suppression of TEAD1 signaling activity. (A) Schematic illustration of the generation of chromaffin cell-specific *TEAD1* overexpression (cTOV) mice by crossing *Pnmt*-Cre mice with conditional *TEAD1* knock-in mice. (B) Plasma epinephrine (EP) levels in cTOV and control (Con) mice following acute cold stimulation. (C) UMAP visualization of single-cell transcriptomic profiles from adrenal medullae of cTOV and Con mice. Chromaffin cells (dashed circle) were further subdivided into three transcriptionally distinct subclusters (clusters 1-3). (D) Feature plot showing the distribution of chromaffin-cell subclusters in cTOV and Con mice. (E) Quantification of chromaffin-cell subcluster abundance in cTOV and Con mice. (F) Monocle pseudotime analysis showing the differentiation trajectory of the three chromaffin-cell subclusters. (G) Monocle pseudotime analysis comparing chromaffin-cell maturation states between cTOV and Con mice. (H) Violin plots showing expression of *Pnmt* and *Th* in chromaffin cells from cTOV and Con mice. (I) Volcano plot showing differentially expressed genes between cTOV and Con chromaffin cells. (J) scMetabolism analysis showing altered metabolic pathways in chromaffin cells from cTOV and Con mice. (K) Western blot analysis of TEAD1, Th, and Pnmt protein expression in adrenal medullary tissues from cTOV and Con mice. (L) Representative immunofluorescence staining of TEAD1 and Pnmt in adrenal medullae from cTOV and Con mice. (M) TEAD1 signaling activity in primary chromaffin cells isolated from cTOV and Con mice. (N) TEAD1 signaling activity following TEAD1 knockdown or overexpression of TEAD family members (TEAD1-4) in PC12 cells. (O) Proposed model illustrating the role of TEAD1 in chromaffin-cell differentiation. Excessive TEAD1 expression promotes early chromaffin-cell differentiation but impairs terminal maturation through suppression of TEAD1 signaling activity. Data are presented as mean ± SEM from three independent experiments (n = 3).

To investigate the underlying mechanism, we performed single-cell RNA sequencing of adrenal medullae from cTOV and control littermates. Chromaffin cells were identified based on Th expression and further resolved into three distinct subpopulations (Fig. 3C). Comparative analysis revealed a marked reduction of cluster 1 and expansion of cluster 2 in cTOV mice, whereas cluster 3 remained largely unchanged (Fig. 3D-E). Monocle trajectory analysis indicated that cluster 1 represented a relatively immature chromaffin-cell state and further demonstrated a global shift toward reduced maturation in cTOV chromaffin cells (Fig. 3F-G).

Consistent with this observation, expression of the terminal differentiation marker Pnmt was significantly decreased in cTOV chromaffin cells, whereas Th expression showed a modest increase (Fig. 3H). The proliferation-associated genes Ccnd1 and Pcna were significantly downregulated in cTOV chromaffin cells (SupFig. 3A). Differential expression analysis revealed upregulation of Th, Slc18a1, and Id1, accompanied by downregulation of key chromaffin maturation genes, including Chga, Chgb, Phox2a, and Pnmt (Fig. 3I). Metabolic pathway analysis further demonstrated significant impairment of tyrosine metabolism, a hallmark pathway required for catecholamine biosynthesis (Fig. 3J).

These transcriptional alterations were validated at the protein level. Western blotting demonstrated robust TEAD1 overexpression accompanied by a marked reduction in Pnmt protein abundance (Fig. 3K). Immunofluorescence staining confirmed efficient TEAD1 overexpression in chromaffin cells and further revealed substantial loss of Pnmt expression throughout the adrenal medulla (Fig. 3L).

Given the inhibitory effect of TEAD1 overexpression on chromaffin-cell maturation, we directly examined TEAD1 signaling activity using an 8×MCAT luciferase-mCherry reporter (SupFig. 3B). Surprisingly, primary chromaffin cells isolated from cTOV mice exhibited significantly reduced reporter activity compared with controls, indicating suppression of endogenous TEAD1 signaling despite elevated TEAD1 expression (Fig. 3M).

To determine whether this phenomenon was cell-autonomous, we performed complementary studies in PC12 cells. Remarkably, both Tead1 knockdown and TEAD1 overexpression resulted in reduced TEAD1 reporter activity (Fig. 3N), suggesting that TEAD1 transcriptional output is not linearly correlated with TEAD1 abundance. Similar observations were subsequently reproduced in pancreatic β cells, pancreatic α cells, and intestinal L cells (SupFig. 3C-E), indicating that this uncoupling between TEAD1 expression and TEAD1 signaling activity may represent a conserved feature of neuroendocrine cells.

To investigate the unique regulatory features of TEAD1 compared with other TEAD family members, we reanalyzed TEAD1 and TEAD4 ChIP-seq datasets generated in HEK293 cells. The top 50 TEAD1– and TEAD4-bound target genes, ranked according to peak-binding scores, were subjected to motif enrichment analysis. We found the TEAD1 binding motif (MCAT) tended to be flanked by more GC pairs than did the TEAD4 binding motif (SupFig. 3F). To validate the hypothesis that TEAD1 transcriptional activity can be affected by sequences near the MCAT motif, we generated 2 luciferase reporters, 3xMCAT-GC and 3xMCAT-AT. In the 3xMCAT-GC reporter, each MCAT motif was flanked by a GC pair, while in 3xMCAT-AT, each MCAT motif was flanked by an AT pair (SupFig. 3G). The luciferase assay revealed that, compared with a flanking GC pair, an AT pair had a stronger repressive effect on TEAD1 or YAP1-TEAD1 pathway activities, as indicated by the results for the 3xMCAT-GC and 3xMCAT-AT reporters (SupFig. 3H). In contrast to our TEAD1 findings, the flanking sequences had no significant effect on TEAD4 pathway activity (Data not shown), suggesting that there are specific and additional mechanisms regulating the transcriptional activity of TEAD1. These findings suggest that the distinct motif-binding preferences of TEAD1 may partially underlie its regulatory specificity compared with other TEAD family members.

Collectively, these findings demonstrate that increased TEAD1 expression does not necessarily enhance TEAD1 signaling. TEAD1 overexpression drives partial differentiation while preventing terminal maturation of chromaffin cells (Fig. 3O). Excessive TEAD1 expression suppresses endogenous TEAD transcriptional activity, resulting in impaired chromaffin-cell maturation and reduced catecholamine production. These data identify TEAD1 signaling activity, rather than TEAD1 abundance itself, as the critical determinant of chromaffin-cell differentiation.

### TEAD reporter-based functional screening identifies HTR5A as a therapeutic target in pheochromocytoma

Given that TEAD1 transcriptional activity more accurately reflects chromaffin-cell differentiation status than TEAD1 expression alone, we next explored whether TEAD1 signaling could be exploited as a functional platform for therapeutic target discovery. Differential expression analysis of tumor chromaffin cells identified 16 significantly enriched G protein-coupled receptors (GPCRs), including HTR5A (Fig. 4A), suggesting potential upstream regulators of TEAD1 signaling.

**Figure 4.**
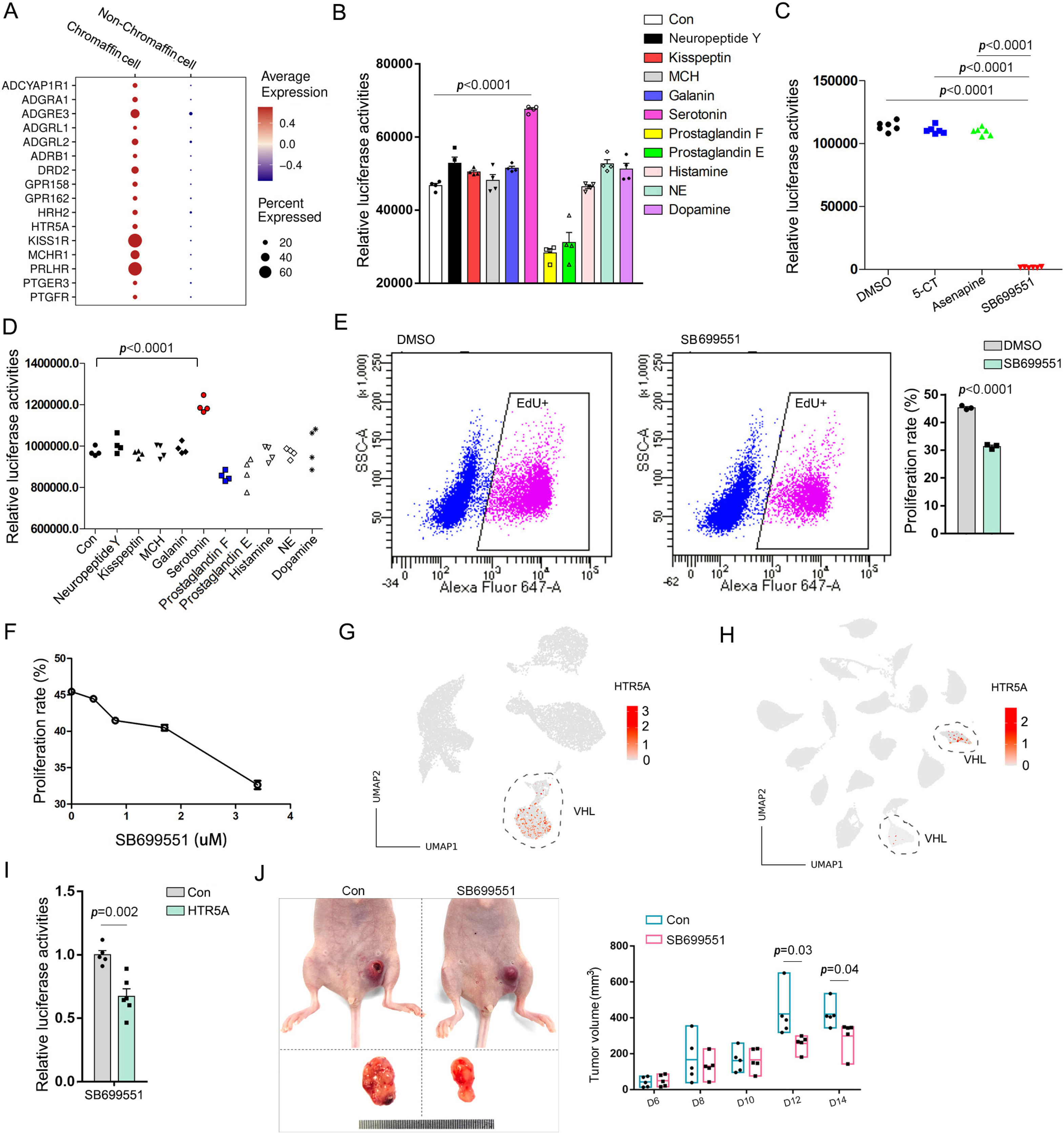
A TEAD1 activity-based screening platform identifies HTR5A as a therapeutic regulator of chromaffin-cell state. (A) Bubble plot showing differential expression of G protein-coupled receptors (GPCRs) between tumor chromaffin cells and non-chromaffin cell populations. (B) TEAD1 signaling reporter (TSR) assay evaluating the effects of selected GPCR ligands on TEAD1 signaling activity in PC12 cells. (C) Effects of HTR5A antagonists on TEAD1 signaling activity measured by TSR. (D) MKI67 reporter assay evaluating the effects of selected GPCR ligands on proliferative activity in PC12 cells. (E) EdU incorporation assay showing the effects of SB699551 on PC12-cell proliferation. (F) Dose-response analysis of SB699551-mediated inhibition of PC12-cell proliferation. (G) Feature plot showing *HTR5A* expression in single-cell transcriptomic datasets. (H) Feature plot showing *HTR5A* expression in the integrated single-nucleus transcriptomic dataset from 18 pheochromocytoma and normal adrenal samples. (I) HTR5A biosensor assay demonstrating inhibition of HTR5A signaling by SB699551. (J) Tumor xenograft experiments showing the in vivo antitumor effects of SB699551. Data are presented as mean ± SEM from three independent experiments (n = 3).

To functionally interrogate these candidates, we employed the TEAD signaling reporter (TSR) system and evaluated ten corresponding ligands in PC12 cells. Among the compounds tested, serotonin markedly increased TSR activity, indicating activation of TEAD1 signaling (Fig. 4B). We subsequently assessed small-molecule antagonists targeting HTR5A and found that SB699551 significantly suppressed TSR activity (Fig. 4C), suggesting that HTR5A signaling contributes to maintenance of endogenous TEAD1 transcriptional output.

Because therapeutic inhibition of pheochromocytoma requires both suppression of tumor growth and modulation of tumor-cell state, we next incorporated a proliferation reporter into our screening strategy. A human MKI67 promoter reporter (−1207 to +316 bp; MR) was generated to quantitatively assess proliferative activity (SupFig 3I-J). Luciferase assays demonstrated that serotonin significantly enhanced MR activity (Fig. 4D). Consistent with these observations, EdU incorporation assays revealed that SB699551 potently inhibited PC12-cell proliferation in a dose-dependent manner (Fig. 4E-F).

Pharmacokinetic analysis by mass spectrometry demonstrated rapid intracellular accumulation of SB699551, reaching peak concentrations approximately 60 minutes after treatment before gradually declining (SupFig. 3K). To identify potential biomarkers for patient stratification, we analyzed both single-cell and single-nucleus RNA sequencing datasets from human pheochromocytoma specimens. These analyses revealed elevated HTR5A expression in a subset of tumors (Fig. 4G-H), suggesting that HTR5A expression may serve as a predictive biomarker for therapeutic responsiveness to SB699551.

To directly monitor HTR5A signaling, we further engineered a NanoLuc-based HTR5A biosensor (SupFig. 3L). Using this system, SB699551 effectively suppressed HTR5A signaling activity (Fig. 4I), confirming on-target pathway inhibition. Finally, therapeutic efficacy was evaluated in a PC12 xenograft model. Treatment with SB699551 resulted in significant inhibition of tumor growth, with tumor volume diverging from control animals after day 12 and progressively decreasing thereafter (Fig. 4J). These findings identify the HTR5A as a previously unrecognized regulator of pheochromocytoma cell proliferation and establish HTR5A inhibition as a potential therapeutic strategy for a molecularly defined subset of pheochromocytomas.

### A subpopulation of human chromaffin tumor cells is sensitive to SB699551

To investigate the biological consequences of HTR5A inhibition, we performed targeted metabolomic profiling in primary human pheochromocytoma cells (PhPCs) following treatment with the HTR5A antagonist SB699551. Metabolic analysis revealed a trend toward reduced levels of vanillylmandelic acid (VMA), the major catecholamine degradation product, whereas the catecholamine precursors tyramine and phenylethylamine were significantly increased (Fig. 5A, SupFig. 4A), indicating substantial remodeling of catecholamine metabolism upon HTR5A inhibition.

**Figure 5.**
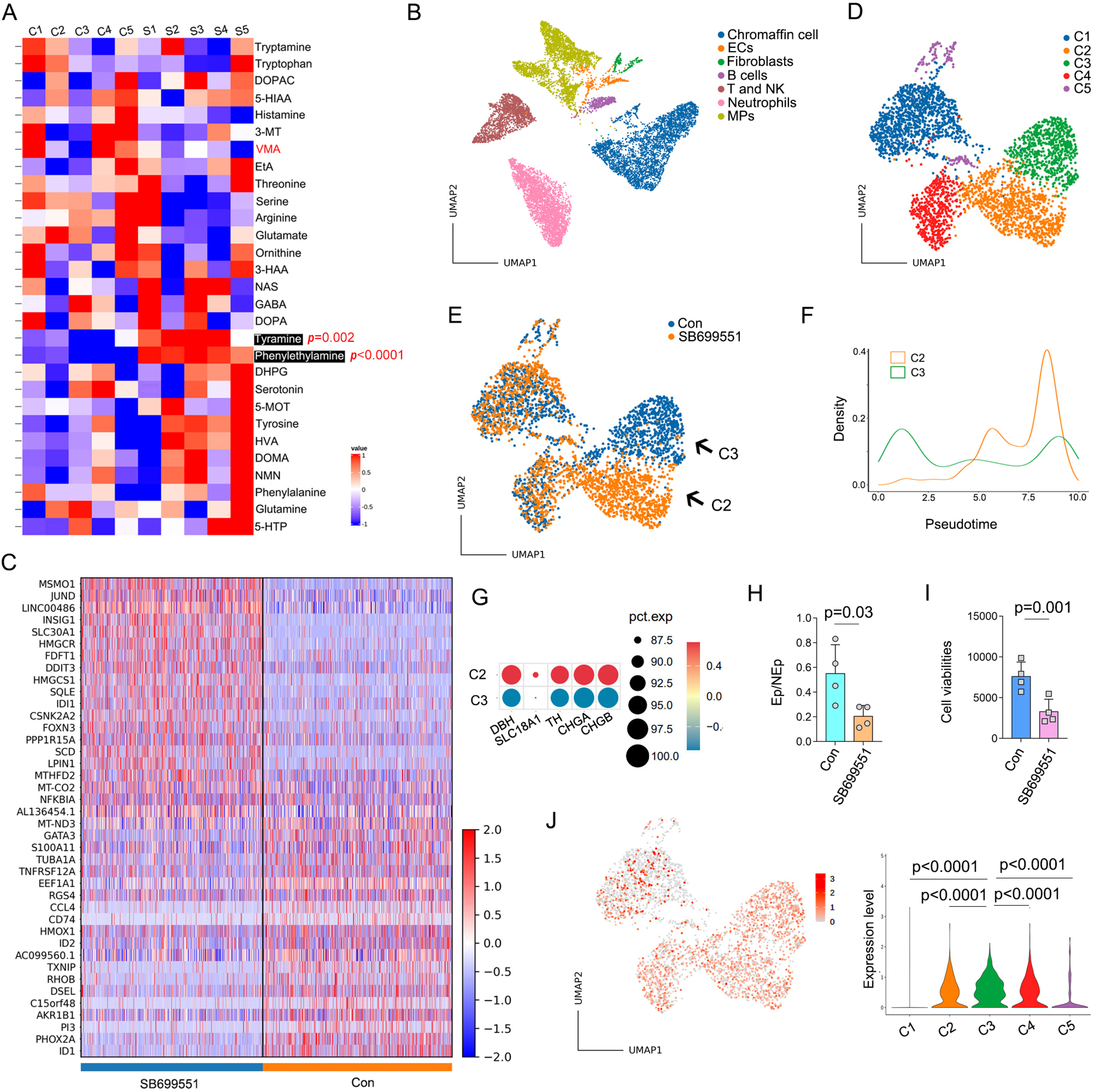
SB699551 partially phenocopies the effects of TEAD1 overexpression in human pheochromocytoma cells. (A) Targeted metabolomic analysis showing metabolite alterations in primary pheochromocytoma cells following SB699551 treatment. (B) UMAP visualization of single-cell transcriptomic profiles from primary pheochromocytoma cells treated with SB699551 or vehicle control (Con). (C) Heatmap showing the top differentially expressed genes in tumor chromaffin cells following SB699551 treatment. (D) UMAP visualization showing reclustering of tumor chromaffin cells into five transcriptionally distinct subpopulations (C1-C5). (E) Changes in chromaffin-cell subpopulation composition following SB699551 treatment. SB699551 treatment resulted in expansion of the C2 population and near-complete depletion of the C3 population. (F) Monocle pseudotime analysis comparing the differentiation states of C2 and C3 chromaffin-cell subpopulations. (G) Bubble plots show increased expression of differentiation-associated markers, including *DBH*, *SLC18A1*, *TH*, *CHGA*, and *CHGB*, in C2 compared with C3. (H) The change of epinephrine-to-norepinephrine (Ep/NEp) ratio following SB699551 treatment in primary pheochromocytoma cells. (I) The change of cell viability following SB699551 treatment in primary pheochromocytoma cells. (J) Feature plots showing TEAD1 expression across chromaffin-cell subpopulations. Violin plots quantify TEAD1 expression levels among individual subpopulations.

To further characterize the cellular response to SB699551, we performed single-cell RNA sequencing of control and SB699551-treated PhPCs. A total of 12,924 cells were profiled and classified into chromaffin cells and six additional cell populations based on established lineage markers (Fig. 5B, SupFig. 4B). Differential expression analysis identified profound transcriptional changes within the chromaffin-cell compartment following SB699551 treatment. Notably, expression of multiple regulators associated with immature neural and neuroendocrine states, including ID1, ID2, PHOX2A, and DCLK1, was significantly reduced (Fig. 5C), suggesting a shift away from progenitor-like transcriptional programs.

To resolve chromaffin-cell heterogeneity, chromaffin cells were further reclustered into five distinct subpopulations (Fig. 5D, SupFig. 4C). SB699551 treatment resulted in striking alterations in subpopulation composition, characterized by near-complete depletion of subcluster 3 and marked expansion of subcluster 2 (Fig. 5E, SupFig. 4D). Pseudotime trajectory analysis demonstrated that cells within subcluster 2 occupied a more differentiated state than those in subcluster 3 (Fig. 5F). Importantly, subcluster 2 displayed enhanced expression of genes associated with a differentiated noradrenergic chromaffin-cell phenotype, including DBH, TH, SLC18A1, CHGA, and CHGB, whereas PNMT expression remained unchanged (Fig. 5G). Consistently, the epinephrine-to-norepinephrine (Ep/NEp) ratio was significantly reduced following SB699551 treatment (Fig. 5H). In addition, SB699551 markedly decreased cell viability (Fig. 5I). Interestingly, TEAD1 expression was highest in subcluster 3, the population most profoundly depleted following SB699551 treatment (Fig. 5J), suggesting that this TEAD1-high chromaffin-cell population may represent a state particularly sensitive to upstream regulation of the TEAD1 pathway.

Collectively, these findings suggest that SB699551 partially phenocopies the effects of TEAD1 overexpression. Specifically, SB699551 promotes the differentiation of TEAD1-high chromaffin tumor cells, as evidenced by increased expression of differentiation-associated markers. However, SB699551 failed to induce PNMT expression or enhance epinephrine biosynthesis, indicating that it does not fully recapitulate terminal adrenergic maturation. In addition, SB699551 significantly reduced cell viability. These results further suggest that TEAD1-high chromaffin tumor cells represent the primary cellular population responsive to SB699551 treatment.

### CXXC5 and L1CAM mediate TEAD1-dependent regulation of chromaffin-cell proliferation and maturation

To identify downstream effectors of the SB699551-TEAD1 signaling axis, we established human induced pluripotent stem cell (iPSC)-derived chromaffin organoids (Fig. 6A). Flow cytometric analysis demonstrated efficient differentiation, with approximately 69% of cells co-expressing TH and PNMT (Fig. 6B), confirming successful generation of chromaffin-like cells.

**Figure 6.**
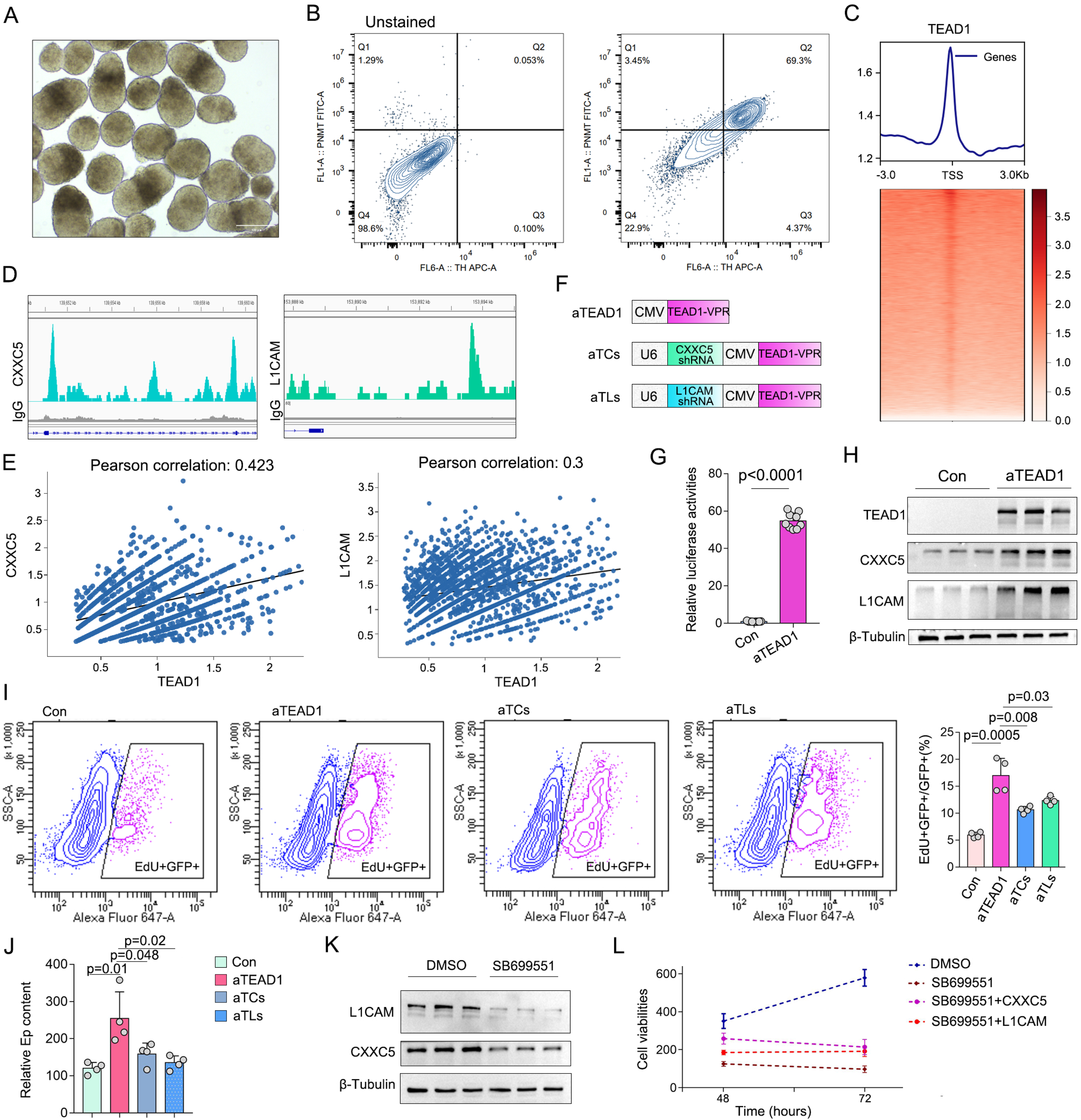
CXXC5 and L1CAM mediate TEAD1-dependent regulation of chromaffin-cell proliferation and maturation. (A) Representative bright-field image of a human iPSC-derived chromaffin organoid. Scale bar, 100 μm. (B) Flow cytometric analysis showing the proportion of TH– and PNMT-double-positive cells in chromaffin organoids. (C) Genome-wide heatmap of TEAD1 occupancy centered on transcription start sites (TSSs) in human chromaffin organoids (±3 kb). (D) Representative TEAD1 binding peaks at the promoter regions of *CXXC5* and *L1CAM*. (E) Correlation analysis between TEAD1 expression and *CXXC5* or *L1CAM* expression in tumor chromaffin cells. (F) Schematic illustration of dominant-positive TEAD1 (aTEAD1) and derivative constructs co-expressing aTEAD1 with *CXXC5* shRNA (aTCs) or *L1CAM* shRNA (aTLs). VPR denotes the VP64-p65-Rta transcriptional activation domain. (G) Luciferase reporter assay showing activation of TEAD1 signaling by aTEAD1. (H) Western blot analysis showing induction of CXXC5 and L1CAM expression following aTEAD1 overexpression in chromaffin organoids. (I) EdU incorporation assay showing the effects of aTEAD1, aTCs, and aTLs on chromaffin-organoid cell proliferation. (J) Effects of aTEAD1, aTCs, and aTLs on epinephrine content in chromaffin organoids. (K) Western blot analysis showing changes in CXXC5 and L1CAM expression following SB699551 treatment in PC12 cells. (L) Cell viability analysis showing the effects of CXXC5 or L1CAM overexpression on SB699551-mediated growth suppression in chromaffin organoids. Data are presented as mean ± SEM from three independent experiments (n = 3).

Using these organoids, we performed ChIP-seq analysis to identify direct TEAD1 target genes (Fig. 6C). Prominent TEAD1 binding peaks were detected within the promoter regions of CXXC5 and L1CAM, suggesting direct transcriptional regulation by TEAD1 (Fig. 6D). Consistent with this observation, single-cell transcriptomic analyses of both human pheochromocytoma and embryonic chromaffin-cell datasets revealed positive correlations between TEAD1 expression and the expression of CXXC5 and L1CAM (Fig. 6E, SupFig. 5A).

To determine whether these genes functionally mediate TEAD1 signaling, we generated a constitutively active TEAD1 construct by fusing TEAD1 to the VPR transcriptional activation domain (aTEAD1). Simultaneously, shRNAs targeting either CXXC5 (aTCs) or L1CAM (aTLs) were incorporated into the same vector, allowing TEAD1 pathway activation and target-gene knockdown within the same cells (Fig. 6F). TEAD1 reporter assays confirmed robust pathway activation by aTEAD1, resulting in an approximately 55-fold increase in TEAD1 reporter activity (Fig. 6G). Western blot analysis further demonstrated marked induction of endogenous CXXC5 and L1CAM expression following aTEAD1 expression (Fig. 6H), supporting their identity as downstream TEAD1 effector genes.

Functional studies revealed that aTEAD1 significantly enhanced chromaffin-organoid proliferation as measured by EdU incorporation. Importantly, knockdown of either CXXC5 or L1CAM partially attenuated the proliferative effects of aTEAD1 (Fig. 6I), indicating that both genes contribute to TEAD1-dependent growth regulation. Similar results were observed for chromaffin-cell function. aTEAD1 significantly increased epinephrine production, whereas silencing of CXXC5 or L1CAM partially diminished this effect (Fig. 6J), suggesting that these factors also participate in TEAD1-mediated chromaffin-cell maturation.

We next examined whether CXXC5 and L1CAM contribute to the pharmacological effects of HTR5A inhibition. In PC12 cells, SB699551 treatment markedly reduced CXXC5 and L1CAM protein abundance (Fig. 6K). Consistent with their role as downstream effectors, overexpression of either CXXC5 or L1CAM, which were validated by western blotting (SupFig. 5B), partially rescued the reduction in organoid viability induced by SB699551 (Fig. 6L). Together, these findings identify CXXC5 and L1CAM as direct TEAD1 target genes that mediate key biological effects of the HTR5A-TEAD1 signaling axis, linking TEAD1 activity to chromaffin-cell proliferation, maturation, and therapeutic responsiveness.

## DISCUSSION

Our study showed that both TEAD1 overexpression and TEAD1 depletion resulted in reduced TEAD1 signaling activity in chromaffin cells, indicating that TEAD1 abundance and TEAD1 pathway output are not linearly correlated. This observation was consistently reproduced in chromaffin cells, pancreatic α cells, β cells, and intestinal L cells, suggesting that such uncoupling may represent a conserved feature of neuroendocrine cells. These findings support a model in which optimal, rather than maximal, TEAD1 abundance is required to maintain effective TEAD1 transcriptional activity.

The mechanism underlying this phenomenon remains incompletely understood. One possible explanation is that excessive TEAD1 occupancy generates non-productive promoter complexes. In our previous study [11], we demonstrated that high levels of TEAD1 binding at promoter regions can interfere with the recruitment of RNA polymerase to chromatin, thereby suppressing transcriptional activation. Based on these observations, we speculate that supraphysiological TEAD1 expression may lead to excessive occupancy of TEAD1-binding elements and impair productive assembly of the transcriptional machinery. Consequently, although TEAD1 abundance is increased, transcriptional output from TEAD1 target genes may be reduced. Under this model, insufficient TEAD1 limits promoter occupancy, whereas excessive TEAD1 restricts efficient RNA polymerase recruitment, resulting in a bell-shaped relationship between TEAD1 abundance and TEAD1 signaling activity.

This model may also help explain the paradoxical phenotype observed in chromaffin-cell–specific TEAD1 overexpression mice. Although elevated TEAD1 expression promoted the transition of immature chromaffin cells toward an intermediate differentiation state, reduced TEAD1 signaling activity impaired acquisition of the fully mature chromaffin-cell phenotype, leading to diminished PNMT expression and epinephrine production. We therefore propose a two-stage model of chromaffin-cell differentiation in which TEAD1 abundance facilitates lineage progression, whereas optimal TEAD1 signaling activity governs terminal maturation and functional identity. Importantly, the observation that both TEAD1 overexpression and depletion impair TEAD1 activity across multiple neuroendocrine cell types raises the possibility that this regulatory principle may extend beyond chromaffin cells and represent a general mechanism controlling neuroendocrine cell maturation.

Beyond the identification of HTR5A as a therapeutically actionable target, an important implication of this study is the establishment of a TEAD activity-based functional screening platform for neuroendocrine tumors. Traditional target-discovery approaches are typically centered on differential gene expression or genetic dependency, which may not adequately capture changes in cellular functional states. In contrast, the TSR system directly reports TEAD1 transcriptional output, a signaling readout that more accurately reflects chromaffin-cell maturation than TEAD1 expression alone. By linking pathway activity to cellular differentiation status, this strategy enables the identification of upstream regulators capable of modulating both tumor growth and neuroendocrine function.

Collectively, our findings suggest that TEAD1 activity, rather than TEAD1 abundance, serves as a functional indicator of neuroendocrine cell state and may provide a broadly applicable framework for therapeutic target discovery across neuroendocrine diseases.

## METHODS

### Compounds and peptides

All compounds and peptides used in this study were purchased from Tocris Bioscience (Bristol, UK) unless otherwise noted. This includes SB699551 (#3188), histamine dihydrochloride (#3545/50), dopamine hydrochloride (#3548/50), serotonin hydrochloride (#3547), galanin (1-29) (#2696), prostaglandin F2α (#4214), prostaglandin E2 (#2296), 5-carboxamidotryptamine maleate (#0458/10), asenapine maleate (#3737/10), noradrenaline bitartrate (#5169/50), neuropeptide Y (#1153), kisspeptin 10 (#4243) and MCH (#3806). For animal injections, SB699551 was purchased from MedChemExpress (#HY-103100). L-DOPA was purchased from Sigma-Aldrich (#D9628; Sigma-Aldrich, St. Louis, MO).

### Clinical samples

Human pheochromocytoma samples were collected from patients and submitted with a requisition form, which included informed consent and patient clinical data. All protocols were approved by the Ethics Committee of Qilu Hospital of Shandong University (Protocol KYLL-202312-054-1). Patient demographics are summarized in Supplementary Table 1.

### Mouse

Pnmt-Cre mice were obtained from GemPharmatech (Nanjing, China). In this strain, a P2A-Cre recombinase cassette was inserted downstream of exon 3 of the endogenous phenylethanolamine N-methyltransferase (*Pnmt*) gene.

For TEAD1 knock-in mouse, the “CAG promoter-loxP-PGK-Neo-6xSV40 pA-loxP-Kozak-Human TEAD1 CDS-WPRE-BGH pA”cassette was cloned into intron 1 of ROSA26.

To generate chromaffin cell-specific TEAD1 overexpression mice (cTOV), Pnmt-Cre mice were crossed with TEAD1 knock-in mice, resulting in selective TEAD1 overexpression in Pnmt-positive adrenal chromaffin cells.

All mice were housed in pathogen-free facilities with a 12h light/dark cycle and had free access to water and food. All mice used for the experiments were males. All animal experiments were conducted in accordance with the Guide for the Care and Use of Laboratory Animals published by the National Institutes of Health.

### Mouse primary chromaffin cell isolation

Excised adrenal glands from 4-month male C57BL/6J mice were placed in 4 °C physiological saline solution (PSS; in mM: 80 Na-glutamate, 55 NaCl, 6 KCl, 1 MgCl_2_, 2 CaCl_2_, 10 HEPES, 10 glucose, pH 7.0, 315 mosmol L−1) and then adipose and capsule were removed. Medullary fragments were rinsed with 5 mL Ca_2_+/Mg_2_+-free PPS and digested (37°C, 10 min) with 1 mL of PPS-Ca_2_+ containing 30 U/mL papain (Sigma-Aldrich, St. Louis, MO; #1.07144) and 0.5 mg/mL BSA (Sigma-Aldrich, #A7906). The enzyme solution was removed and replaced with 1 mL of PPS-Ca_2_+ containing 3 U/mL collagenase F (Sigma-Aldrich, #C7926), 0.5 mg/mL BSA, and 100 μM CaCl_2_ (37 °C, 10 min). Isolated chromaffin cells from 5 cTOV and 5 Control mice were subject to single-cell sequencing. All mouse experiments performed were approved by the Institutional Animal Care and Use Committee (Protocol DWU-2024-089).

### Cell lines

Rat PC-derived PC12 cells, and alpha TC1 cells were purchased from ATCC (#CRL-1721.1, # CRL-2934), and cultured and maintained according to the protocol from ATCC. GLUTag and Min6 cells were purchased from Sigma (#SCC623, #SCC652).

### Differentiation of hiPSCs into chromaffin-like cells

Chromaffin-like cells were generated from human induced pluripotent stem cells (hiPSCs) using a modified stepwise differentiation protocol adapted from previously described methods for axial progenitor, trunk neural crest, sympathoadrenal progenitor, and adrenal chromaffin cell differentiation [13–14].

Briefly, hiPSCs were dissociated into single cells and seeded onto Matrigel-coated plates at a density of approximately 5.5 × 10^4^ cells/cm² in axial progenitor induction medium containing 4 μM CHIR99021 and 20 ng/mL FGF2. Y-27632 (10 μM) was included during the first 24 h after plating. Cells were maintained under these conditions for 3 days.

For trunk neural crest induction, day 3 cells were dissociated and replated onto Matrigel-coated plates at approximately 3.0 × 10^4^ cells/cm². Cells were cultured in DMEM/F12 supplemented with N2, MEM non-essential amino acids, GlutaMAX, 2 μM SB431542, 1 μM CHIR99021, 1 μM DMH1, and 15 ng/mL BMP4. Y-27632 (10 μM) was added only during the first 24 h after replating. To enhance Schwann cell precursor–associated signaling, recombinant human NRG1 (10 ng/mL) was supplemented from day 6 to day 9.

On day 9, cells were transitioned to sympathoadrenal progenitor induction medium consisting of DMEM/F12 supplemented with B27, N2, MEM non-essential amino acids, and GlutaMAX, together with 50 ng/mL BMP4, 50 ng/mL recombinant SHH C24II, and 1.5 μM purmorphamine. Cells were maintained in this medium until day 14.

For chromaffin maturation, day 14 cells were dissociated and transferred to ultra-low-attachment plates to generate suspension aggregates. Aggregates were cultured in DMEM/F12 supplemented with GlutaMAX, 2% B27, 0.5% BSA, 1× penicillin-streptomycin, 5 ng/mL BMP4, 10 μM dexamethasone, and 100 nM phorbol 12-myristate 13-acetate (PMA). Half of the medium was replaced every 2 days with fresh maturation medium. Cells were maintained under these conditions for an additional 6-9 days before downstream analyses.

### Human primary pheochromocytoma cell isolation and culture

Fresh PC-derived tissue was dissected from patients’ tumors. Tissue was subsequently minced into small pieces, digested in Hank’s Buffer Salt Solution (HBSS) supplemented with collagenase type 4 (2.5 mg/mL; Sigma-Aldrich, #C4-28) and deoxyribonuclease I (0.05 mg/mL; Sigma-Aldrich, #10104159001) for 1 hour in a 37℃ shaking incubator at 150 rpm. Digested tissue was dispersed into a single-cell suspension by gentle pipetting, followed by filtration through a 100 μm mesh filter, and pelleting by centrifugation (1000 rpm, 1 min). The cell pellet was resuspended in DMEM supplemented with 15% fetal bovine serum (FBS) and penicillin-streptomycin, and subsequently maintained as a suspension of nonadherent single cells in Falcon 6-well culture plates (#351146; Corning, Corning, NY). In this medium, fibroblasts attached, while the neuroendocrine cells remained in suspension. After recovery for 4-6 hours, subsets of primary PC cells were treated with either 4 μM SB699551 or DMSO as the vehicle control for 16-24 hours and then prepared for single-cell RNA sequencing.

### Tumor xenograft experiments

Tumor xenograft studies employed 4-6-week-old female pathogen-free BALB/c nude mice (BALB/c NCR-Nu; GemPharmatech, Nanjing, China) that were congenitally athymic (n=5). 2 x 10^6^ PC12 cells were transplanted subcutaneously in the inguinal region. SB699551 was dissolved in DMSO first and then diluted with corn oil for in vivo injection; DMSO-corn oil was used as control. The injection of SB699551 and control started on day 5 post-transplantation. Mouse weights and tumor sizes were monitored every 2 days. Tumor volumes were calculated using the formula: Vol = L × W^2^ × 0.5. All mouse experiments performed were approved by the Institutional Animal Care and Use Committee (Protocol DWU-2024-089).

### Luciferase reporters and assay

Reporter construction. To generate luciferase reporters of expression, the following sequences were cloned from the respective gene promoter regions and inserted into the adenoviral backbone to drive NanoLuc luciferase and P2A mCherry expression: human MKI67 between −1207 and +316. 8xMCAT along with firefly luciferase were generated by PCR of relevant sequences in the Hop-flash construct (Addgene, #83467) and inserted into the adenoviral backbone to generate the TEAD1 signaling reporter (TSR).

Luciferase assay. Luciferase assays were performed with a Promega Dual-Luciferase® Kit (#E1960; Promega Corporation, Fitchburg, WI) according to the manufacturer’s instructions. Renilla luciferase activities or total DNA content served as internal controls. All plasmids utilized in this study were validated by Plasmid-EZ (Genwiz, Waltham, MA).

### Plasmids

Plasmids containing human TEAD1 (#OHu18002), CXXC5 (#OHu31350), and L1CAM (#OHu27607) open reading frames (ORFs) were purchased from GenScript (Piscataway, NJ). The respective gene coding sequences (CDS) were subcloned into the adenoviral plasmid backbone and fused with both a FLAG tag and P2A GFP. Cleaved GFP was used as a marker of adenoviral transduction efficiency.

### shRNA Knockdown

The target sequence for human CXXC5 is GAAAGACTGGCCATCAGATTT and for human L1CAM is CCACTTGTTTAAGGAGAGGAT. A scrambled RNA were used as a negative control, and the sequence of hairpin is: CCTAAGGTTAAGTCGCCCTCGCTCGAGCGAGGGCGACTTAACCTTAGG.

### Cell viability measurement

Cell viability was determined using the RealTime-Glo™ MT Cell Viability Assay (Promega, #G9711), which was performed according to the manufacturer’s instructions.

### Adenovirus packaging

In this study, TEAD1, L1CAM, and CXXC5 constructs were packaged into Ad5 adenoviral vectors. Briefly, HEK293 cells were seeded in 10 cm dishes to reach 50-70% confluence and transfected with the adenoviral vectors using Lipofectamine™ 3000. Cells were incubated at 37 °C for 5-8 h, after which the transfection medium was replaced with complete growth medium. Cells and supernatants were harvested 7-14 days post-transfection and subjected to three freeze-thaw cycles to lyse cells and release viral particles. Cell debris was removed by centrifugation at 7,000 × g for 10 min at 4 °C, and the resulting supernatant was collected as the primary viral stock. For viral amplification, fresh HEK293 cells were infected with the crude viral stock to increase viral titer, which was measured using the Adeno-X™ Rapid Titer Kit (Takara Bio, #632250).

### Cold exposure test

To evaluate stress-induced epinephrine secretion, mice were subjected to acute cold exposure. Briefly, mice were individually housed in a 4°C cold chamber for 2 h with free access to water. Blood samples were collected after cold exposure. To prevent epinephrine oxidation, sodium metabisulfite was added to the collected blood samples at a final concentration of 0.1%. Plasma was isolated by centrifugation and stored at −80°C until analysis. Plasma epinephrine concentrations were quantified using an Epinephrine ELISA Kit (Elabscience, #E-EL-0045).

### Immunostaining

Immunostaining was performed as previously described [15]. The following primary antibodies were used: anti-TEAD1 rabbit monoclonal (1:100; #ab133533; Abcam, Cambridge, UK), anti-TEAD1 rabbit monoclonal (1:100; #12292; Cell Signaling Technology, Danvers, MA), anti-TH mouse monoclonal (1:300; #45648; Cell Signaling Technology), anti-CHGA mouse monoclonal (1:100; #36468; Cell Signaling Technology), and anti-PNMT rabbit polyclonal (1:300; #13217-1-AP; Proteintech). Secondary antibodies included Alexa Fluor™ 488 chicken anti-rabbit IgG (H+L) secondary antibody (#A-21441, Thermo Fisher Scientific, Waltham, MA) and Alexa Fluor™ 594 chicken anti-mouse IgG (H+L) secondary antibody (#A-21201, Thermo Fisher Scientific). The images were captured by a Nikon confocal A1R microscope with 40x objective (Nikon, Tokyo, Japan).

### Western blotting

Western blotting was performed as previously described [16]. The following primary antibodies were used: anti-TEAD1 rabbit monoclonal (#12292; Cell Signaling Technology), anti-FLAG rabbit monoclonal (#14793; Cell Signaling Technology), anti-GAPDH mouse monoclonal (#97166; Cell Signaling Technology), anti-TH mouse monoclonal (#45648; Cell Signaling Technology), and anti-PNMT rabbit polyclonal (#13217-1-AP; Proteintech). IRDye® 800CW donkey anti-rabbit IgG (#926-32213) and IRDye® 680RD (#926-68070) goat anti-mouse IgG were used as secondary antibodies (LI-COR Biotechnology, Lincoln, NE). Western blots were imaged using an Odyssey Clx imaging system (LI-COR Biotechnology).

### EdU incorporation assay

EdU (5-ethynyl-2’-deoxyuridine) incorporation was detected using Click-iT Plus EdU flow cytometry assay kits (#C10646, #C10634; Thermo Fisher) according to the manufacturer’s instructions. Briefly, cells were seeded and allowed to attach overnight, followed by treatment with the indicated compounds for 48 h. During the final 1 hour of treatment, cells were incubated in FBS-free tissue culture medium containing 10 μM EdU. Cells were then fixed with 4% formaldehyde for 15 min, permeabilized for 15 min, incubated with 0.5 mL Click-iT® Plus reaction cocktail for 30 min, rinsed, and analyzed using a BD FACSAria™ flow cytometer (BD Biosciences, Franklin Lakes, NJ). Detection of GFP fluorescence from the GFP reporter expressed by the adenoviral constructs served as a marker of adenovirus-transduced cells. The proliferation rate was calculated as the percentage of EdU/GFP double-positive cells among GFP-positive cells (EdU⁺GFP⁺/GFP⁺).

### HTR5A biosensor construction

Design of the intramolecular HTR5A biosensors was based on the split NanoLuc luminescence assay. At the 5’ end of the human HTR5A CDS, a cleavable signal sequence was included to promote membrane localization. Additionally, a V2 vasopressin receptor tail polypeptide (V2 tail) was added to the C terminus to promote β-arrestin2 recruitment followed by the N-terminal fragment of NanoLuc luciferase (N-NanoLuc). The constructs then included a cleavable T2A element followed by the C-terminal fragment of NanoLuc fused to human ARRB2 (β-arrestin2) at both its N– and C-termini.

### Targeted metabolomics

High performance liquid chromatography-mass spectrometry (HPLC-MS). Metabolomic analysis and SB699551 measurement were performed by a Sciex 5500 Q-Trap mass spectrometer equipped with an electrospray ionization (ESI) source (Sciex, Framingham, MA). The mass spectrometer was coupled with an Agilent 1290 infinity UHPLC system (Agilent, Santa Clara, CA). Mobile phase A consisted of 25 mM ammonium formate and 0.1% formic acid (FA); mobile phase B was composed of acetonitrile containing 0.1% FA. Samples were placed in the 4℃ automatic injector with the column temperature set to 45℃. An injection volume of 2 μL was applied to all samples. A gradient (90% B at 0 min, 40% B at 18 min, 90% B at 18.1 min, 90% B at 18.1-23 min) was then initiated at a flow rate of 300 μL/min. Quality control (QC) samples were set for each interval of experimental samples in the sample queue to detect and evaluate the stability and repeatability of the system. The source conditions of ESI were as follows: source temperature 450℃, ion source gas1: 60, ion source gas2: 60, curtain gas: 30, ion spray voltage floating 5000 V, MRM mode was used to detect ion pairs.

#### SB699551 measurement

PC12 cells were seeded in 6-well plates (106 cells per well) and cultured for 24 hours. After SB699551 treatment (1 μM) in different time points, PC cells were dissociated with trypsin (#25200056, Thermo Fisher) and washed three times with PBS. Intracellular SB699551 content was measured in cell lysates via HPLC-MS according to the methods above. SB699551 purchased from Tocris Bioscience (#3188, Bristol, UK) was used as a standard.

#### Data processing

MultiQuant software v3.0 (Sciex) was used to extract chromatographic peak area and retention time. QC samples were processed together with the biological samples. Metabolites in the QC samples with coefficient of variation (CV) less than 30 % were denoted as reproducible measurements.

#### Data analysis

Statistical significance (p-value < 0.05) was determined using a two-tailed t-test. Metabolites with p value <0.05, fold change >1.5 or fold change <0.67 were considered as significantly changed metabolites. For hierarchical clustering, Cluster3.0 (http://bonsai.hgc.jp/∼mdehoon/software/cluster/software.htm) and the Java Treeview software (http://jtreeview.sourceforge.net) were used. A Euclidean distance algorithm for measures of similarity and an average linkage clustering algorithm (which uses clustering based on the centroids of the observations) were selected.

### Single-cell RNA sequencing

Initial analysis of raw sequencing data. Raw reads were processed to generate gene expression profiles using CeleScope v1.15.0 (Singleron Biotechnologies, Köln, Germany). Briefly, barcodes and unique molecular identifiers (UMIs) were extracted from R1 reads and corrected. Adapter sequences and poly-A tails were trimmed from R2 reads. Trimmed reads were aligned against the GRCm38 transcriptome using STAR RNA-seq aligner software v2.6.1b (https://github.com/alexdobin/STAR/releases/tag/2.6.1b). Uniquely mapped reads were assigned to genes using FeatureCounts software v2.0.1 (https://subread.sourceforge.net/featureCounts.html). Successfully assigned reads with the same cell barcode, UMI, and gene were grouped together to generate gene expression matrices for further analysis.

Quality control, dimension-reduction, and clustering. Under Python 3.9, Scanpy software v1.8.2 (https://scanpy.readthedocs.io/en/stable/) was used for quality control, dimensionality reduction, and clustering. For each sample dataset, the expression matrix was filtered using the following exclusion criteria: 1) cells with a gene count < 200 or with a top 2% gene count; 2) cells with a top 2% UMI count; 3) cells with mitochondrial content > 5%; and 4) genes expressed < 5 cells. After filtering, 89,021 cells were retained for downstream analyses, with an average of 2,026 genes and 13,683 UMIs per cell. The raw count matrix was normalized by total counts per cell and logarithmically transformed into normalized data matrix. The top 2,000 variable genes were selected by setting flavor as ‘seurat’ for guided clustering. Principle Component Analysis (PCA) was performed on the scaled variable gene matrix, and the top 20 principal components were used for clustering and dimensional reduction. Cells were separated into 25 clusters using the Louvain algorithm and setting resolution parameters at 1.2. Cell clusters were visualized by Uniform Manifold Approximation and Projection (UMAP).

Batch Effect removal. Batch effects among samples were removed via Harmony software v1.0 using the top 20 principal components from the PCA results.

Differentially expressed gene (DEG) analysis. To identify differentially expressed genes (DEGs), the scanpy.tl.rank_genes_groups() function was employed based on Wilcoxon rank sum test with default parameters. DEGs were selected according to the genes expressed in > 10% of cells in either of the compared groups of cells and with an average log(Fold Change) value > 1. The Benjamini-Hochberg procedure was used to calculate the adjusted p values with p < 0.05 considered statistically significant.

Cell type annotation. Cell-type recognition was performed by Cell-ID, which is a multivariate approach that extracts gene signatures from each individual cell and performs cell identity recognition using hypergeometric tests (HGT). Dimensionality reduction was performed on normalized gene expression matrices through multiple correspondence analysis, where both cells and genes were projected in the same low dimensional space. A gene ranking was then calculated for each cell to obtain the most featured genesets of that cell. HGT were performed on these genesets against the SynEcoSys database, which contains featured genes of all the cell types. For cluster annotation, the frequency of every cell-type was calculated in each cluster, and the highest frequency cell type was chosen as the identity of the cluster.

Transcription factor regulatory network analysis (pySCENIC). A transcription factor network was constructed by pyscenic (v0.11.0) using scRNA expression matrix and transcription factors in AnimalTFDB. First, GRNBoost2 predicted a regulatory network based on the co-expression of regulators and targets. CisTarget was then applied to exclude indirect targets and to search transcription factor binding motifs. After that, AUCell was used for regulon activity quantification for every cell. Cluster-specific TF regulons were identified according to Regulon Specificity Scores (RSS) and the activity of these TF regulons were visualized in heatmaps.

### Trajectory analysis

Cell trajectory analysis was performed using the Monocle2 package (v2.24.0) in R. Highly variable genes identified from the Seurat object were used to construct the CellDataSet. Cells were ordered in pseudotime using the DDRTree dimensionality reduction algorithm followed by the orderCells function. The trajectory was rooted in sympathoadrenal progenitor cells according to the expression of established developmental markers. Genes with dynamic expression patterns along pseudotime were identified for downstream analysis.

### ChIP-seq

Chromatin immunoprecipitation (ChIP) experiments. ChIP was performed as previously described17. Briefly, Dynabeads Protein A beads (#10001D; Life Technologies Corporation, Carlsbad, CA) were placed in a total volume of 25 μL and washed twice with 200 μL ice-cold 140 mM RIPA buffer (10 mM Tris-HCl, 140 mM NaCl, 1 mM EDTA, 0.5 mM EGTA, 0.1% SDS, 0.1% Na-deoxycholate, 1% Triton X-100, 1 mM PMSF, 1x proteinase inhibitor Cocktail, 20 mM Na-butyrate, pH 7.5). This was followed by resuspension in RIPA buffer to a final volume of 200 μL. A total volume of 5 µL anti-TEAD1 antibody (#ab133533, Abcam) or IgG control was added to the bead suspension, and followed by incubation on a tube rotator for at least 2.5 hours at 4°C. hPheo1 cells were fixed in 1% formaldehyde for 10 min and quenched with 125 mM glycine for 5 min. Cells were incubated in 150 μL lysis buffer (50 mM Tris-HCl pH 8.0, 10 mM EDTA pH8.0, 0.5% SDS, 1 mM PMSF, 1x proteinase inhibitor cocktail, 20 mM Na-butyrate) for 20 min on ice, then sonicated and centrifuged. The supernatant was transferred to a 1 mL tube containing suspended antibody-coated Protein A beads, followed by incubation on a tube rotator overnight at 4°C. Next, the beads were transferred to a new tube and incubated in 100 μL ChIP elution buffer (10 mM Tris-HCl, 5 mM EDTA, 300 mM NaCl, 0.5% SDS, pH 8.0) containing 5 µL proteinase K (Qiagen, 20 mg/ml stock) at 55°C for 2 hours and then at 65°C for 4 hours. The eluate was transferred to a fresh tube, and the enriched DNA was purified by phenol-chloroform, followed by dissolution in 50 μL TE buffer. A NEB Next Ultra II DNA Library Prep Kit for Illumina (#E7645S; New England Biolabs, Ipswich, MA) was used for library construction according to manufacturer’s instructions. The libraries were sequenced on a Hiseq X-ten instrument (Illumina) set for paired-end 150 base pair sequencing.

ChIP-seq analysis. Clean reads were mapped to the human reference genome using Bowtie2 software v2.54 (https://bowtie-bio.sourceforge.net/bowtie2/index.shtml). Reads from mitochondrial DNA, reads with mapping quality <30, and PCR-duplicated reads were all removed. High-quality mapping reads were subjected to further peak calling. Macs2 software was used to call peaks with a p-value < 0.05. Peaks were annotated by the ChIPseeker software package. The findMotifsGenome.pl tool within the HOMER software package v4.11 (http://homer.ucsd.edu/homer/motif/) was used for Motif analysis. Input files consisted of the peak file and the genome fasta file. The DNA sequence was extracted according to the peak file, and the sequence was compared with the Motif database to obtain the Motif.

### Public ChIP-seq, scRNA-seq and snRNA-seq data analysis

The original snRNA-seq matrix files for 18 human PC and healthy adrenal tissues were previously described [17]. The matrix files for human fetal adrenal tissues were obtained from previously described data deposited in the Gene Expression Omnibus (GEO) (GSE157329, GSE195929, GSE147821) [18–20]. Joint analysis was performed for post-batch effect removal. Processed BED and WIG files were downloaded from ChIP-atlas (http://ChIP-atlas.org/) [21]. WIG files were opened in Integrative Genomics Viewer (IGV) (https://igv.org/doc/desktop/) to obtain the “PEAK” graphs. CLC Genomics Workbench 12 (Qiagen) was used to extract the target gene information on “PEAK score”, annotation, and nearby gene information. Specifically, TEAD1 and TEAD4 ChIP-seq in HEK293 cells were employed to perform various target gene analyses. TEAD1 and TEAD4 binding motif graphs were generated using WebLogo 3 (http://weblogo.berkeley.edu/logo.cgi).

### Statistical Analysis

All the data were presented as the mean ± SEM. Differences between groups were evaluated using unpaired Student’s t-test or ANOVA (one-way and two-way) followed by Tukey’s post hoc tests using GraphPad Prism software (version 8; GraphPad Software, Boston, MA. p < 0.05 was considered statistically significant.

### Data Availability

ScRNA-seq and ChiIP-seq data have been deposited at GEO and will be publicly available as of the date of publication.

### Author contributions

L.X., chromaffin organoid induction. X.L., immunostaining. F. Y., animal work. J.Q., molecular experiments. Y.Z., animal experiments. M.C., collected clinical samples and isolated primary cells. R.L., V.K.Y., R.W.T., K.Z., and L.C. contributed to data analyses. F.L., X.H., and F.L. conceived and supervised the work and wrote the manuscript with input from the co-authors. F.L. designed the project.

